# Signal Inhibitory Receptor on Leukocytes-1 recognizes S100 proteins

**DOI:** 10.1101/2021.05.14.444196

**Authors:** Matevž Rumpret, Helen J. von Richthofen, Maarten van der Linden, Geertje H. A. Westerlaken, Cami Talavera Ormeño, Teck Y. Low, Huib Ovaa, Linde Meyaard

## Abstract

Signal inhibitory receptor on leukocytes-1 (SIRL-1) is an inhibitory receptor with a hitherto unknown ligand, and is expressed on human monocytes and neutrophils. SIRL-1 inhibits myeloid effector functions such as reactive oxygen species (ROS) production. We here identify S100 proteins as SIRL-1 ligands. S100 proteins are composed of two calcium-binding domains. Various S100 proteins are damage-associated molecular patterns (DAMPs) released from damaged cells, after which they initiate inflammation by ligating activating receptors on immune cells. We now show that the inhibitory SIRL-1 recognizes individual calcium-binding domains of all tested S100 proteins. Blocking SIRL-1 on human neutrophils enhanced S100 protein S100A6-induced ROS production, showing that S100A6 suppresses neutrophil ROS production via SIRL-1. We conclude that SIRL-1 is an inhibitory receptor recognizing the S100 protein family of DAMPs.

## Introduction

Inhibitory immune receptors set thresholds for immune cell activation and ensure prompt cessation of immune responses, preventing immunopathology [1]. Signal inhibitory receptor on leukocytes-1 (SIRL-1, VSTM1) is an inhibitory receptor without known ligands, expressed on human granulocytes and monocytes [2]. It dampens Fc-receptor-mediated reactive oxygen species (ROS) production in neutrophils and monocytes and neutrophil extracellular trap formation *in vitro* [3–5].

S100 proteins are small proteins with member-specific functions [6]. The human genome encodes twenty-one S100s [7, 8]. Although functionally diverse, S100s share a conserved structure comprising two Ca^2+^-binding EF-hand domains, one S100 protein-specific and the other common to other Ca^2+^-binding proteins. Each EF-hand domain is composed of two α-helices connected by a short Ca^2+^-binding loop, giving rise to a helix–loop–helix motif, and the two EF-hand domains are joined by a short hinge region to form a full S100 protein [9]. Some S100s are damage-associated molecular patterns (DAMPs) released from damaged cells [10, 11]. Additionally, S100s such as S100A8 and S100A9 are also actively secreted from activated immune cells. After release into extracellular space, many S100s interact with pattern recognition receptors such as TLR4 and RAGE and promote inflammation [12–14]. Here, we show that some S100s engage SIRL-1 to suppress S100-induced ROS in neutrophils.

## Materials and Methods

### Construction of 2B4 NFAT-GFP reporter cells

2B4 NFAT–GFP cell line is a T cell hybridoma cell line in which GFP expression is controlled by the transcriptional factor NFAT. LAIR-1–CD3ζ construction has been described [15], and SIRL-1–CD3ζ construction was performed similarly, yielding a 269 AA chimera comprising the SIRL-1 signal peptide and ectodomain (MTAEFLSLLC…APSMKTDQFK) fused to the CD3ζ transmembrane and cytoplasmic domains (LCYLLDGILF…ALHMQALPPR). The construct was transduced into a 2B4 NFAT–GFP cell line, in which ligation of the chimera by an antibody or a ligand results in NFAT promoter-driven GFP expression [16]. Three days after transduction, cells were sorted for high SIRL-1 surface expression and subcloned by limiting dilution. We selected one clone with high surface SIRL-1 expression and high GFP expression levels after SIRL-1 ligation (“2B4 NFAT–GFP reporter cell assay” below) for further experiments. All cells were maintained in RPMI 1640 with 10% heat-inactivated fetal bovine serum (FBS) and 50 U/ml penicillin–streptomycin (culture medium) unless stated otherwise.

### Peptide synthesis

EF-hand domains of S100s A6, A8, A9, and A12 were synthesized as described [17], with some modifications. Peptide synthesis was performed on a Syro II Multisyntech automated synthesizer with a TIP module. We applied standard 9-fluoronylmethoxycarbonyl (Fmoc) based solid-phase peptide chemistry at a 2 μmol scale, using fourfold excess of amino acids relative to pre-loaded Fmoc amino acid wang-type resin (0.2 mmol/g Rapp Polymere) [17]. For all cycles, double couplings in NMP for 25 min using PyBOP (4 equiv) and DiPEA (8 equiv) were performed. After each coupling, the resin was washed five times with NMP. In between cycles, Fmoc removal was performed with 20% piperidine in NMP for 2×2 min and 1×5 min. After Fmoc removal, the resin was washed three times with NMP. Resins were then washed with ethoxyethane and dried on air. Peptides were cleaved from resins and deprotected with TFA/H_2_O/phenol/iPr_3_SiH (90.5/5/2.5/2 v/v/v/v) for 3 h. After resines were washed with 3×100 μL TFA/H_2_O/phenol/iPr_3_SiH (90.5/5/2.5/2 v/v/v/v), peptides were precipitated with cold Et_2_O/n-pentane 3:1 v/v. Precipitated peptides were washed 3× with Et_2_O, dissolved in H_2_O/CH_3_CN/HOAc (75/24/1 v/v/v), and lyophilized [17].

### 2B4 NFAT–GFP reporter cell assay

We assessed SIRL-1 activation using SIRL-1–CD3ζ 2B4 NFAT–GFP T cell hybridoma reporter cells with wt and LAIR-1–CD3ζ 2B4 NFAT–GFP cells as controls [15]. MaxiSorp Nunc (Figures 1 and 3) or Greiner (Figure 2) 96-well flat-bottom plates were coated overnight at 4°C with 10 μM S100 proteins (Prospec) or fragments (own production), or 5 μg/ml collagen I (Sigma), unless stated otherwise. Mouse-anti-SIRL-1 mAb (clone 1A5, own production; 10 μg/ml), mouse-anti-LAIR-1 mAb (clone 8A8, own production; 10 μg/ml), and Armenian hamster-anti-mouse-CD3 mAb (clone 145-2C11, BD; 10 μg/ml) were coated as controls. After washing the wells with PBS, we seeded 0.5×10^5^ cells/well in the culture medium and incubated them overnight at 37°C. Where indicated, cells were pre-incubated with anti-SIRL-1 F(ab’)_2_ or control F(ab’)_2_ for 30 min before seeding. For anti-CD3 control in reporter assays with pre-incubation with F(ab’)_2_, 1 μg/ml anti-mouse-CD3 was coated to the plate. The following day, GFP expression was measured by flow cytometry (LSR Fortessa; BD Bioscience) following the “Guidelines for the use of flow cytometry and cell sorting in immunological studies” [18]. Results were analyzed with FlowJo 10.0.7r2.

**Figure 1:**
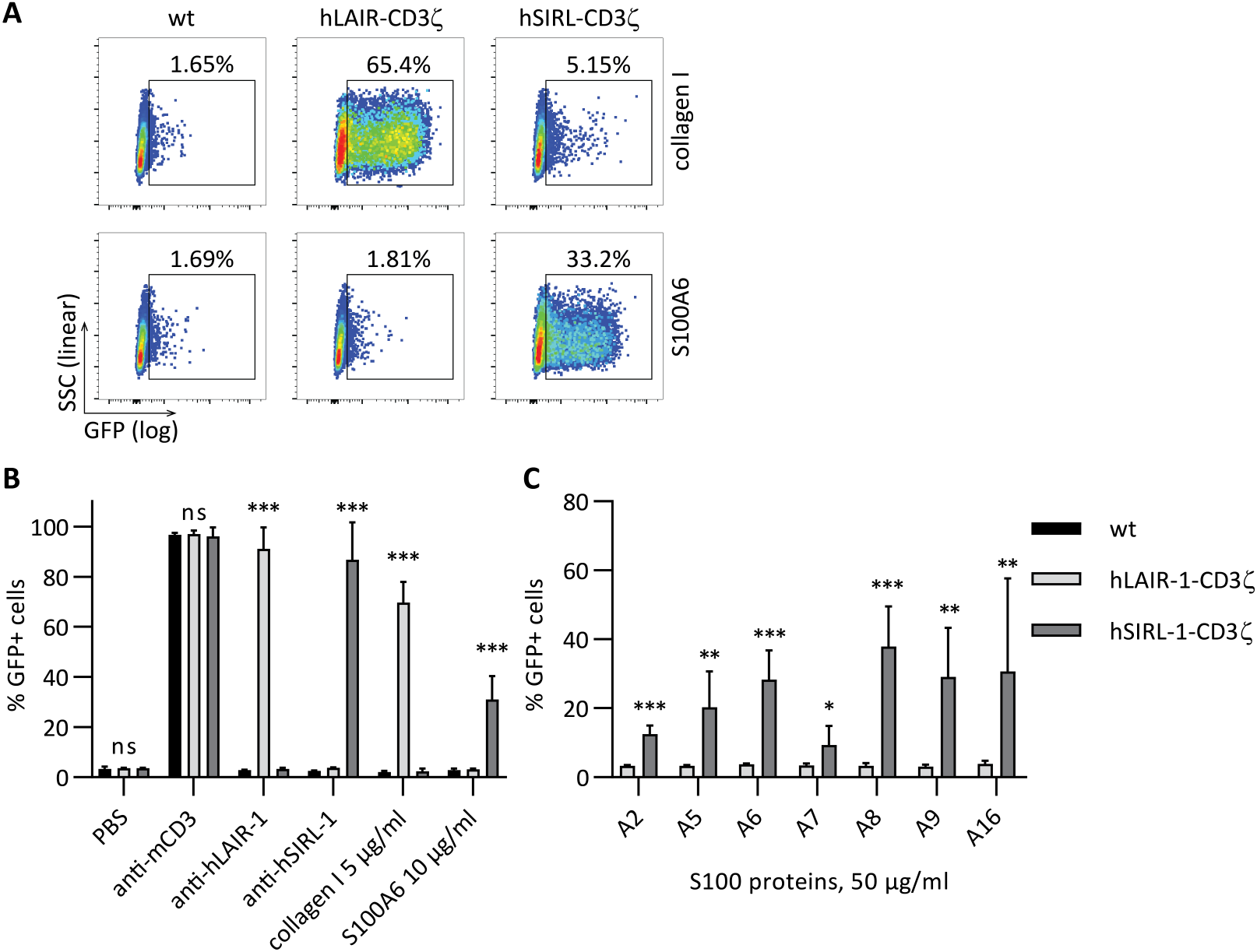
S100 proteins activate SIRL-1 reporter cells. Reporter cells were incubated with antibodies, collagen I, and S100s coated to a MaxiSorp 96-well plate. We assessed receptor activation by measuring GFP expression by flow cytometry. A) Representative dot plots and gating examples are shown. B) Reporter cells stimulated with anti-CD3 mAb, anti-hLAIR-1 mAb, collagen I (a known LAIR-1 ligand), anti-hSIRL-1 mAb and S100A6. C) Reporter cells stimulated with different S100 proteins. Mean and SD of three independent experiments are displayed. Student’s *t*-test with the Holm-Šidák multiple comparison correction. * p<0.05; ** p<0.01; *** p<0.001; ns = not significant. Significance levels for comparison hLAIR-1–CD3ζ to hSIRL-1–CD3ζ are shown.

### Neutrophil isolation and ROS assay

We collected all samples after obtaining informed consent and with the approval of the Medical Research Ethics Committee Utrecht. Neutrophils were isolated from peripheral blood of healthy donors by Ficoll gradient centrifugation [19], with modifications. After erythrocyte lysis, neutrophils were washed with RPMI 1640 with 2% FBS. Neutrophils were then washed and suspended in HEPES buffer (20 mM HEPES, 132 mM NaCl, 6 mM KCl, 1 mM CaCl_2_, 1 mM MgSO_4_, 1.2 mM KH_2_PO_4_, 5 mM D-glucose and 0.5% (w/v) bovine serum albumin, pH 7.4) with 4% FBS. Isolated neutrophils were incubated for 30 minutes at room temperature with 20 μg/ml control F(ab’)_2_ (Southern Biotech) or mouse-anti-SIRL-1 F(ab’)_2_ (own production). We generatead F(ab’)_2_ fragments using Pierce™ Mouse IgG1 Fab and F(ab’)_2_ Preparation Kit (ThermoFisher Scientific #44980). ROS production upon stimulation with or without 10 μM peptide S100A6-1 coated to a Microfluor^®^ 2 white-bottom 96-well plate (ThermoFisher) was determined by AmplexRed assay [3]. Fluorescence was measured in a 96-well plate reader (Clariostar, BMG Labtech) every two minutes for 150 minutes (λ Ex/Em = 526.5-97 / 650-100 nm). We corrected the measurement for spontaneous ROS production by subtracting the signal of PBS-treated cells.

### Statistical analysis

Student’s *t*-test with the Holm-Šidák multiple comparison correction, or paired Student’s *t*-test were performed as indicated. P-values lower than 0.05 were considered statistically significant (* p<0.05; ** p<0.01; *** p<0.001). Statistical analysis was performed with GraphPad Prism 8.

## Results and discussion

### S100 proteins activate SIRL-1

To identify SIRL-1 ligands, we performed pull-downs with recombinant SIRL-1 and LAIR-1 ectodomains from human monocyte and mouse RAW cell line lysates (supplementary information). We identified enriched proteins by mass spectrometry (MS, data available in jPOSTrepo). We used LAIR-1, an inhibitory receptor for collagen [15], as a specificity control. Human collagen II was enriched in LAIR-1 pull-downs. No proteins were significantly enriched in SIRL-1 pull-downs, but we did detect several S100 proteins in increased quantity. Despite S100s commonly appearing as non-specific hits in such experiments, we validated them further.

We tested multiple S100s as ligands for SIRL-1 in a reporter assay, in which ligation of SIRL-1–CD3ζ or LAIR-1–CD3ζ chimera by an antibody or a ligand results in GFP expression (Figure 1). SIRL-1– CD3ζ cells selectively responded to plate-coated anti-SIRL-1 mAb and S100A6 (Figure 1A shows selected scatter plots, and Figure 1B shows quantification). We next investigated additional members of the S100 protein family—A2, A5–A9, and A16—for SIRL-1 activation. All selectively activated the SIRL-1 reporter (Figure 1C), indicating SIRL-1 recognizes S100s. To investigate which S100 protein domains activate SIRL-1, we synthesized peptides spanning individual EF-hand domains of S100s A6, A8, A9, and A12. A sequence alignment [20] indicating EF-hand domains is shown in Figure 2A, and peptide sequences are listed in supplementary table 1. We tested these fragments (Figure 2B) and found that one EF-hand domain was sufficient for SIRL-1 reporter cell activation and that both canonical and S100-specific EF-hand domains of selected S100s engaged SIRL-1. Such broad recognition of ligands is reminiscent of LAIR-1 recognizing different types of extracellular and membrane collagens [21].

**Figure 2:**
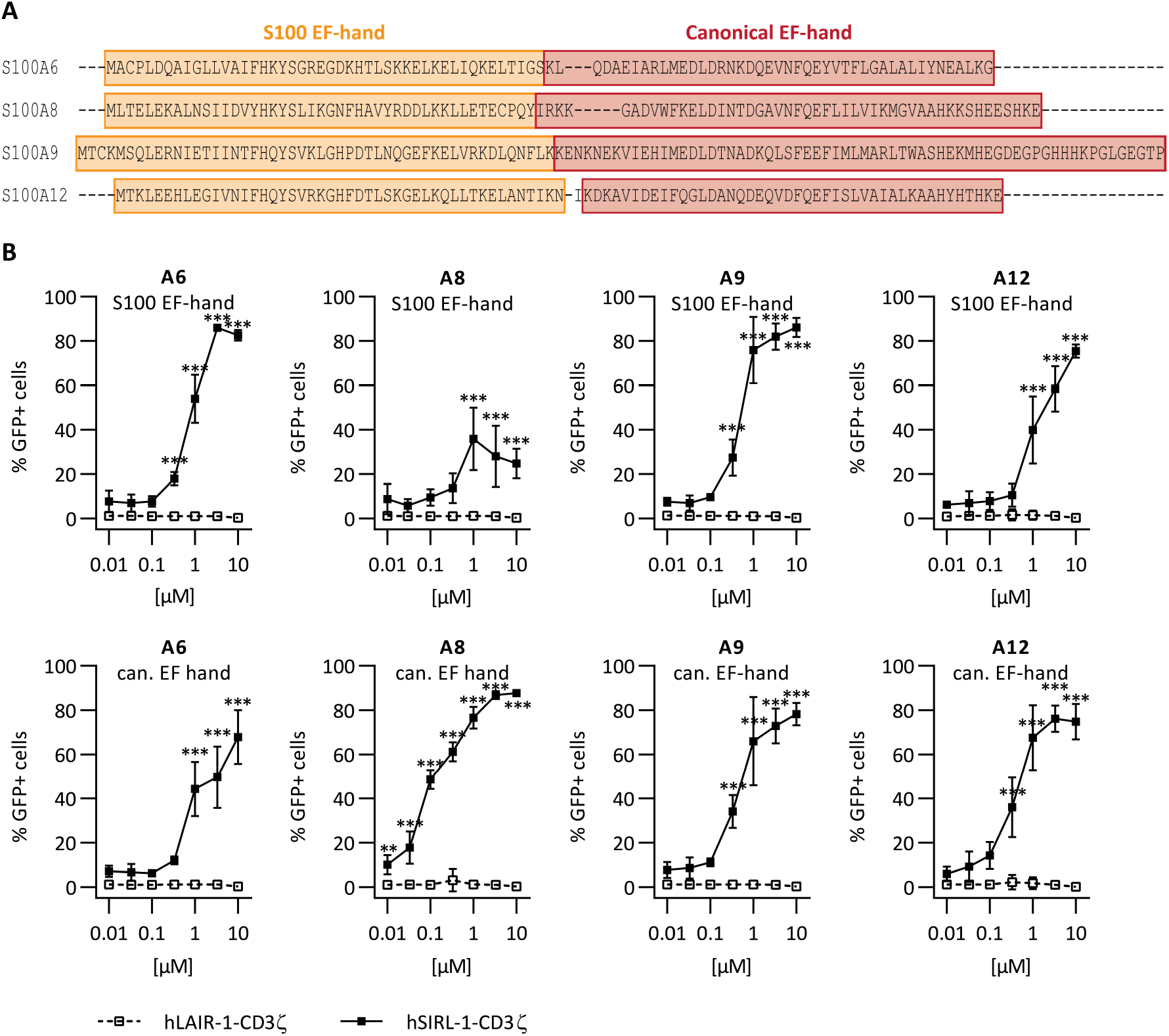
EF-hand domains of S100 proteins activate SIRL-1. A) Sequence alignment of selected S100s. S100-specific (orange) and canonical (red) EF-hand domains are indicated. B) Reporter cells were incubated with EF-hand domains of S100A6, S100A8, S100A9, and S100A12 coated to Greiner 96-well plates. We assessed receptor activation by measuring GFP expression by flow cytometry. Mean and SD of three independent experiments are displayed. Student’s *t*-test with the Holm–Šidák multiple comparison correction. * p<0.05; ** p<0.01; *** p<0.001.

### Anti-SIRL-1 mAb blocks activation of SIRL-1 by S100 proteins

We next performed the reporter assay in the presence of anti-SIRL-1 F(ab’)_2_ or control F(ab’)_2_ (Figure 3A). We used anti-SIRL-1 F(ab’)_2_ instead of the full-length mAb to prevent Fc-receptor engagement, in the experiments where we block SIRL-1. Anti-SIRL-1 F(ab’)_2_, but not control F(ab’)_2_, concentration-dependently blocked S100-induced SIRL-1 reporter cell activation. Thus, S100 protein EF-hand domain recognition by SIRL-1 depends on SIRL-1 ectodomain accessibility. In the case of S100A9, the blockade was not significant, possibly due to the low GFP signal. Upon anti-CD3 stimulation, we observed no blockade by anti-SIRL-1 F(ab’)_2_, excluding steric hindrance by F(ab’)_2_ as the reason for anti-SIRL-1 F(ab’)_2_-specific blockade of S100-induced GFP (Figure 3A).

**Figure 3:**
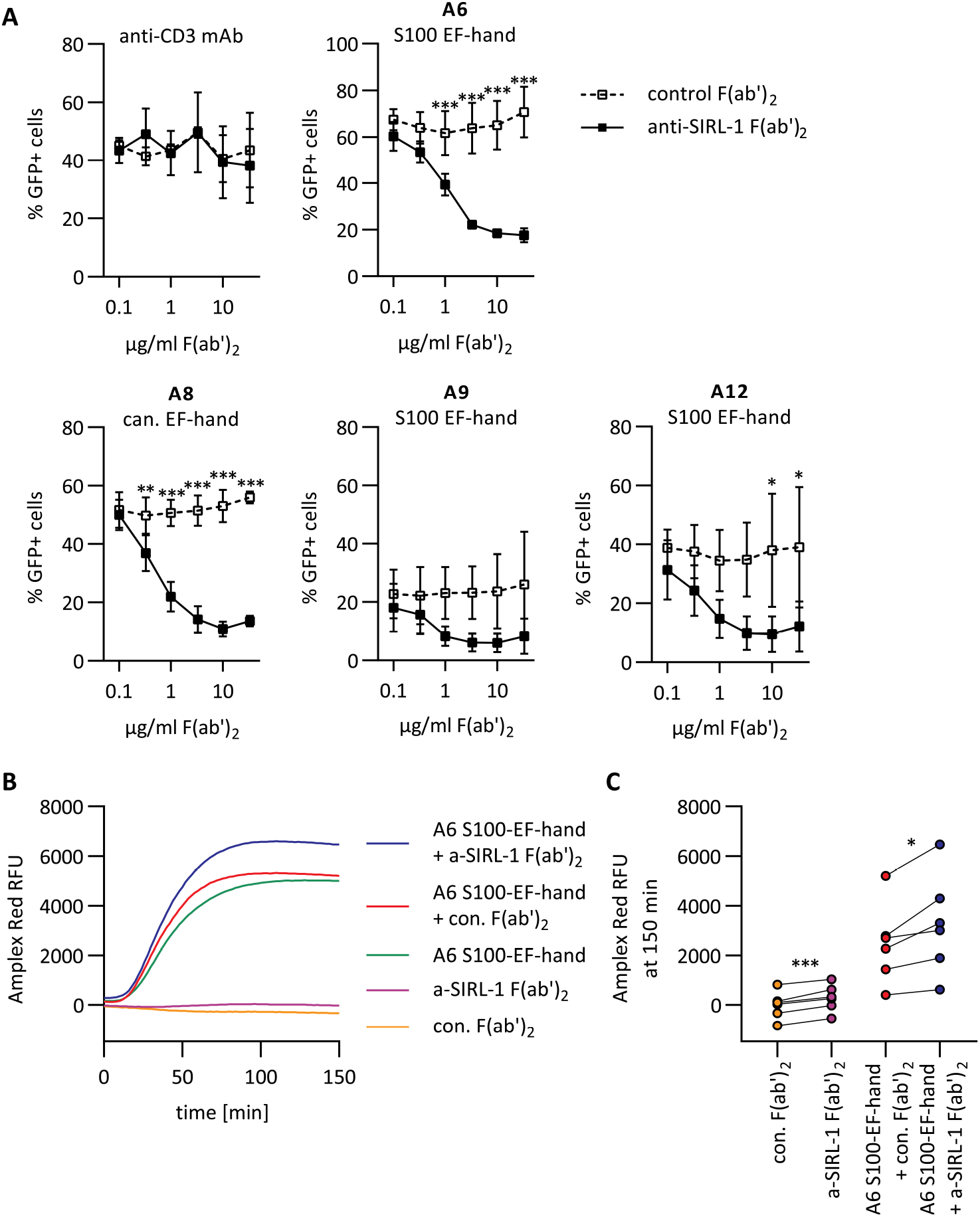
SIRL-1 blockade with anti-SIRL-1 F(ab’)_2_ enhances S100A6-induced ROS in human neutrophils. A) SIRL-1 reporter cells were pre-incubated with varying concentrations of anti-SIRL-1 F(ab’)_2_ or a control F(ab’)_2_. Subsequently, they were incubated with S100 EF-hand domains of S100A6, S100A9, and S100A12, and the canonical EF-hand domain of S100A8 that had been coated to a MaxiSorp 96-well plate. 1 μg/ml coated anti-CD3 mAb was used as a control. We assessed receptor activation by measuring GFP expression by flow cytometry. Mean and SD of three independent experiments are displayed. Student’s *t*-test with the Holm-Šidák multiple comparison correction. B, C) Human neutrophils were pre-incubated with anti-SIRL-1 F(ab’)_2_ or a control F(ab’)_2_ and subsequently stimulated with S100-EF-hand of S100A6 coated to Microfluor 2 96-well plate. ROS production was monitored for 150 minutes. B) ROS production over time, shown for one representative donor. C) ROS production at 150 min for all six donors. Three independent experiments with neutrophils isolated from blood of six healthy donors are shown. Paired Student’s *t*-test. * p<0.05; ** p<0.01; *** p<0.001.

### SIRL-1 blockade enhances S100-instigated ROS production in human neutrophils

SIRL-1 is a negative regulator of FcR-induced ROS production [3], and some S100s induce ROS in neutrophils [22]. Thus, we investigated whether SIRL-1 inhibits S100-induced ROS production. We stimulated freshly isolated human neutrophils with SIRL-1-activating EF-hand domains of S100A5, S100A6, S100A8, S100A9, and S100A12. Of these, only the S100-specific EF-hand of S100A6 induced neutrophil ROS production (Supplementary Figure 1). Upon neutrophil stimulation with the S100-specific EF-hand of S100A6, anti-SIRL-1 F(ab’)_2_ pre-incubated neutrophils exhibited significantly higher ROS production than control F(ab’)_2_ pre-incubated neutrophils (Figure 3B, C), showing SIRL-1 dampens S100A6-induced ROS production in neutrophils. We observed a slight increase in ROS production upon SIRL-1 blockage in the absence of S100A6. This could indicate the presence of S100s or other SIRL-1 ligands in medium with serum, of which recognition was also blocked by anti-SIRL-1 F(ab’)_2_. In agreement with this, theNFAT–GFP SIRL-1–CD3ζ reporter cells typically show slightly higher background GFP levels than wt and LAIR-1–CD3ζ reporter cells.

We could not show a direct interaction between S100s and SIRL-1 (Supplementary figures 2 and 3), even though we initially identified S100s by SIRL-1 pull-down from cell lysates. Additional molecules could be needed for SIRL-1–S100 interaction, which is not uncommon—for instance, TLR4 binds LPS in complex with MD2 and CD14 [23]. Alternatively, the SIRL-1–S100 interaction affinity might be too low for detection in a purified system. Lastly, S100s frequently appear as non-specific hits in MS experiments. The initial SIRL-1 pull-down of S100s might thus have been a chance finding, of which follow-up experiments showed that S100s specifically activate SIRL-1.

### Concluding Remarks

DAMPs can deliver activating as well as inhibitory signals to immune cells—the inhibitory receptor Siglec-10 recognizes high mobility group box 1, a prototypical DAMP, in complex with CD24 [24]. The S100 DAMPs behave similarly: S100A9 binds the inhibitory receptor LILRB1 (CD85j) to modulate NK cell activity [25], and here we show that S100A6 induces SIRL-1-mediated inhibition. What is the benefit of SIRL-1 mediated negative regulation through DAMPs like S100s? When neutrophils first infiltrate the affected tissue to deploy effector mechanisms, the tissue has not been damaged by inflammatory processes and is devoid of DAMPS like S100s. Later in inflammation, effector cells will induce tissue damage and enhance DAMP release. We have previously shown that activated neutrophils gradually downregulate SIRL-1 expression [3]. Incoming neutrophils, however, express high levels of SIRL-1, which can detect S100s and induce an inhibitory signal. This functions as negative feedback for further deployment of tissue-damaging effector mechanisms and reduces the chances of developing immunopathology. In conclusion, we identify the first known SIRL-1 ligands. We demonstrate that S100 proteins activate SIRL-1 and that blocking SIRL-1 with anti-SIRL-1 F(ab’)_2_ fragments enhances S100A6-induced ROS production in neutrophils. Similarly, S100s could modulate monocyte function through SIRL-1, which can be explored in the future. We propose that SIRL-1 broadly recognizes a yet unidentified feature of S100 proteins to help limit tissue damage by immune cells.

## Abbreviations

SIRL-1: Signal inhibitory receptor on leukocytes-1
LAIR-1: Leukocyte-associated immunoglobulin-like receptor-1
ROS: reactive oxygen species;
DAMP: damage-associated molecular pattern;
MS: mass spectrometry;
FBS: fetal bovine serum;
Fmoc: 9-fluoronylmethoxycarbonyl

## Data availability statement

The data supporting the findings of this study are available within this article, in the supplementary material of this article, or are openly available in jPOSTrepo (Japan ProteOme STandard Repository) at: https://repository.jpostdb.org/preview/19411839915f8d78ad99ab9 with the access key: 2145.

## Ethics approval statement for human studies

We collected samples after obtaining informed consent and with the approval of the Medical Research Ethics Committee Utrecht.

## Author contributions

L.M., M.R., and M.v.d.L. conceptualized the study. M.R., H.v.R., M.v.d.L., and G.H.A.W. performed the experiments and analyzed the data. C.T.O. synthesized the peptides. T.Y.L. and M.v.d.L. performed the analysis of the MS experiments. M.R. wrote the initial draft. All authors except H.O. contributed to the reviewing and editing of the final manuscript. H.O. and L.M. supervised the study. L.M. acquired funding for the study.

## Acknowledgments

We want to thank Wim de Lau for help with cloning and expression of the SIRL-1 and LAIR-1 HA-Flag fusion proteins, Albert Heck for help and advice on performing the MS experiments, Lieneke Jongeneel for generation of F(ab’)_2_ fragments, and Florianne Hafkamp for help with performing the GFP reporter assays.

This work was supported by a Vici grant from the Netherlands Organization for Scientific Research awarded to L.M. (NWO, grant no. 91815608).

