## Supplementary information for "Signal Inhibitory Receptor on Leukocytes-1 recognizes S100 proteins"

### Supplementary materials and methods

| | |
|---|---|
| S100A5_1 | METPLEKALTTMVTTFHKYSGREGSKLTLSRKELKELIKKELCLGE |
| S100A5_2 | MKESSIDDLMKSLDKNSDQEIDFKEYSVFLTMLCMAYNDFFLEDNK |
| S100A6_1 | MACPLDQAIGLLVAIFHKYSGREGDKHTLSKKELKELIQKELTIGS |
| S100A6_2 | KLQDAEIARLMEDLDRNKDQEVNFQEYVTFLGALALIYNEALKG |
| S100A8_1 | MLTELEKALNSIIDVYHKYSLIKGNFHAVYRDDLKKLLETECPQY |
| S100A8_2 | IRKKGADVWFKELDINTDGAVNFQEFLILVIKMGVAAHKKSHEESHKE |
| S100A9_1 | MTCKMSQLERNIETIINTFHQYSVKLGHPDTLNQGEFKELVRKDLQNFLK |
| S100A9_2 | KENKNEKVIEHIMEDLDTNADKQLSFEEFIMLMARLTWASHEKMHEGDEGPGHHHKPGLGEGTP |
| S100A12_1 | TKLEEHLEGIVNIFHQYSVRKGHFDTLSKGELKQLLTKELANTIKN |
| S100A12_2 | KDKAVIDEIFQGLDANQDEQVDFQEFISLVAIALKAAHYHTHKE |

**Supplementary table 1: List of S100 fragment peptides.**

EF-hand domains: 1 = S100-specific, 2 = canonical

*Recombinant protein cloning, expression, and purification*

SIRL-1 and LAIR-1 ectodomains (containing their native signal sequences) were PCR-amplified and cloned into pJet2.blunt. Inserts were cut out from the plasmid using PCR primer-encoded restriction enzymes and ligated into pcDNA3.1 expression vector (ThermoFisher) in frame with C-terminal 2×HA and 2×Flag tags. HEK293T cells were transiently (PEI)-transfected with newly generated plasmids. After one day, cells were transferred from RPMI 1640 supplemented with 10% (v/v) heat-inactivated fetal bovine serum and 50 U/ml penicillin–streptomycin to serum-free OptiMEM (ThermoFisher) and incubated for five days at 37°C and 5% $CO_2$. Afterward, the supernatant was collected by centrifugation, and the recombinant proteins were purified by incubation of the clarified supernatant with anti-FLAG (M2) agarose on a rotatory incubator at 4°C overnight. The next day, anti-FLAG (M2) agarose loaded with the SIRL-1-(2×HA-2×Flag) or LAIR-1-(2×HA-2×Flag) ectodomains was washed five times with PBS, and pull-down experiments were performed.

*Monocyte isolation*

All samples were collected after obtaining informed consent and with the approval of the Medical Research Ethics Committee Utrecht. Human monocytes were isolated from the freshly drawn peripheral blood of healthy donors. Whole blood was centrifuged over a Ficoll (GE Healthcare) gradient, and isolated PBMCs were further centrifuged over a Percoll (GE Healthcare) gradient. Purified monocytes were then lysed immediately, as described below.

*Cell lysis, pull-down assays, and mass spectrometry*

Isolated monocytes and mouse RAW cells were lysed by incubation in 50 mM Tris pH 8, 150 mM NaCl, 1% NP-40, and protease inhibitor cocktail (Roche) at 4°C for 30 min. Following lysis, the lysate was cleared by centrifugation at 12,000 g and 4°C for 20 min. Cleared lysates (protein content around 1 mg/ml) were incubated with SIRL-1-HA-Flag or LAIR-1-HA-Flag recombinant proteins bound to anti-FLAG agarose (Sigma) at 4°C overnight. The next day, anti-FLAG agarose was washed six times with PBS, and proteins were eluted with 100 µg/ml FLAG peptide (Sigma). The eluate was then incubated with anti-HA agarose (Sigma) at 4°C overnight. The next day, anti-HA agarose was washed six times with PBS, protease digestion was performed, and samples were submitted to mass spectrometric analysis [1].

*Cell-based binding assay*

Recombinant S100 proteins, anti-SIRL-1, and anti-LAIR-1 antibodies were dissolved in PBS to a final concentration of 10 µg/ml. Collagen I was dissolved in PBS with 2 mM acetic acid to a final concentration of 25 µg/ml. 150 µl of the solutions were incubated in wells of a MaxiSorp 96-well plate at 4°C overnight to allow immobilization. The next day, wells were washed three times with PBS and incubated with 1% (w/v) BSA solution in PBS for 60 min at RT, and the plate was again washed three times with PBS. K562 is a human lymphoblast cell line, which we transduced with full-length SIRL-1 analogous to the LAIR-1-overexpressing K562 cells, described previously [2]. K562 cells (wt, LAIR-1-overexpressing, and SIRL-1-overexpressing) were concentrated to $5x10^6$ cells/ml and incubated with the fluorescent dye calcein dissolved in PBS for 30 min at 37°C in a cell culture incubator. Cells were then washed three times with RPMI 1640 + 1% fetal bovine serum (FBS), and $1.5x10^5$ cells in 100 µl RPMI 1640 + 1% FBS were added to the microtiter plate and incubated for 6 h at 37°C in a cell culture incubator to allow adherence to the plate-coated proteins. After incubation, cells were washed 15 times with RPMI 1640 + 1% FBS, and calcein fluorescence was recorded (Ex/Em = 485/527 nm) in a plate reader before every wash step to measure the decrease in calcein fluorescence intensity as a consequence of cell detachment. Results are shown in supplementary figure 2.

*ELISA-based binding assay*

Recombinant S100 proteins were dissolved in PBS to a final concentration of 10 μg/ml. Anti-SIRL-1 (clone 1A5, own production) and anti-LAIR-1 (clone 8A8, own production) antibodies and bovine serum albumin (BSA) were dissolved in PBS to a final concentration of 5 μg/ml. 100 μl of the solutions were incubated in wells of a MaxiSorp 96-well plate at 4°C overnight to allow immobilization. The next day, wells were washed three times with PBS and incubated with 3% (w/v) BSA solution in PBS for 60 min at RT. The plate was then washed five times with PBS. Fusion proteins containing ectodomains of SIRL-1 or LAIR-1 and the Fc dimerization tag, respectively SIRL-1-Fc or LAIR-1-Fc, were added to the wells to a concentration of 10 μg/ml in PBS + 1% BSA and incubated for 2 h at RT. Wells were washed five times with PBS. Anti-human-IgG-HPR Ab was added to 0,2 μg/ml in PBS + 1% BSA and incubated for 1 h at 4°C. Wells were washed five times with PBS, and ELISA was developed with TMB substrate and stopped with 1 M $H_2SO_4$. Absorbance was measured at 450 nm. Results are shown in supplementary figure 3.

### Supplementary figures

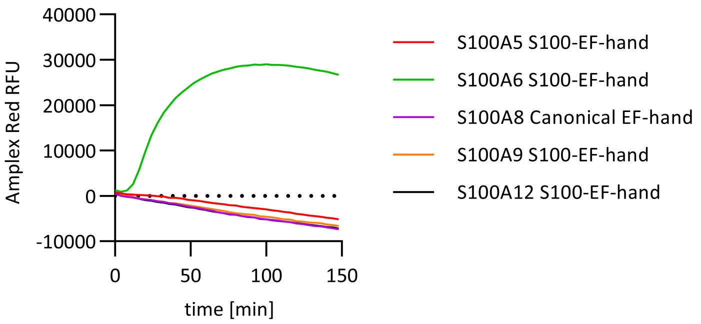


**Supplementary figure 1:** EF-hand motifs of selected S100 proteins were tested for induction of ROS in human neutrophils. The S100-specific EF-hand of S100 protein A6 was the only one that induced ROS. Representative traces of one experiment.

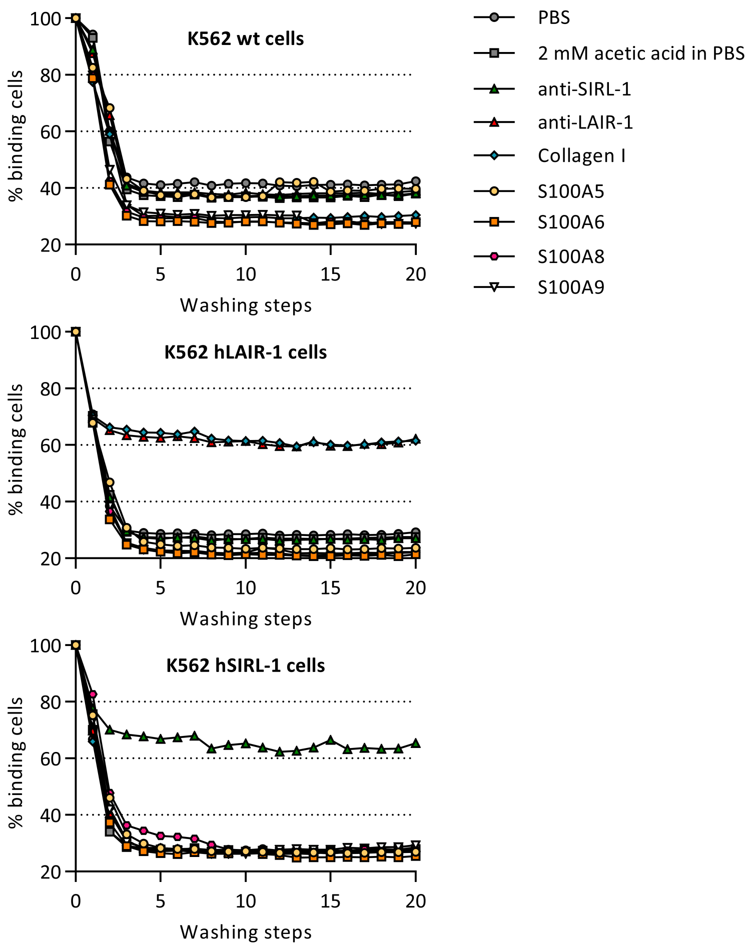


**Supplementary figure 2:** SIRL-1 overexpressing and control K562 cells were incubated with plate-coated S100 proteins as well as control antibodies and ligands. LAIR-1 overexpressing cells bound plate-coated collagen I and anti-LAIR-1 antibody. SIRL-1 overexpressing cells bound to plate-coated anti-SIRL-1 antibody, but no binding was observed to S100 proteins A5, A6, A8, and A9. A representative experiment is shown.

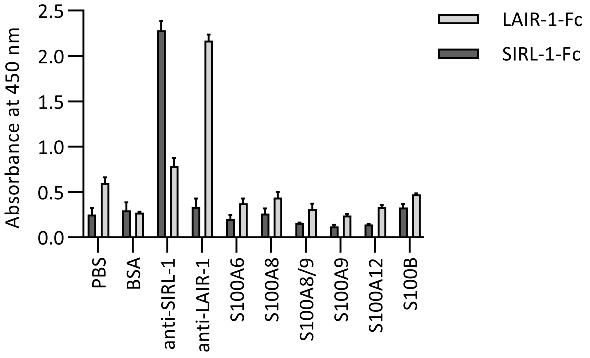


**Supplementary figure 3:** Recombinant SIRL-1-Fc fusion protein selectively bound to anti-SIRL-1, but not to plate-coated S100 proteins A6, A8, A8/9 heterodimer, A9, A12, or B. Recombinant LAIR-1-Fc fusion protein selectively bound to anti-LAIR-1, and was used as a control. Mean and SEM of one representative experiment are shown.
